# Advances in the Design and Functionality of a Compact Multi-Reflecting Time-of-Flight Mass Spectrometer

**DOI:** 10.64898/2026.06.16.732645

**Authors:** Jason Wildgoose, Samantha Ferries, Lee A. Gethings, Matthew E. Daly, Martin E. Palmer, Richard Lock, Johannes P.C. Vissers, James I. Langridge

**Affiliations:** Waters Corporation, Altrincham Road, Wilmslow, SK9 4AX, UK

**Keywords:** multi-reflecting time-of-flight (MRT), high resolution mass spectrometry, omics

## Abstract

**Rationale:** High-resolution mass spectrometry is routinely used for the analysis of complex samples in pharmaceutical, environmental, and omics related studies. Such applications demand instrumentation to be capable of combining sub-ppm mass accuracy, high resolving power, rapid full m/z range acquisition, over a wide dynamic range.

**Methods:** Achieving the above requirements places constraints on analyzer design and performance. Multi-reflecting time-of-flight (MRT) based analyzers have previously been reported as a means of significantly extending the effective flight path in compact TOF designs. Here, further instrument and functionality advances in a compact MRT mass spectrometer design are described.

**Results:** The impact of these enhancements was assessed for targeted and non-targeted omics applications, examining the impact of acquisition speed on resolving power, dynamic range including limits of quantitation, and quantitative precision.

**Conclusion:** The results obtained characterize the performance of the enhanced design features of a compact MRT mass spectrometer. Operation at elevated acquisition rates up to 200 Hz was observed without loss in resolving power, isotopic ratio accuracy, or quantitative precision.

## Introduction

The performance characteristics of mass spectrometers (MS) are largely determined by analyzer design, which influences mass resolving power, mass measurement accuracy, acquisition speed, transmission efficiency, and dynamic range ^1–6^. In recent years, ongoing increases in sample complexity and desired analytical throughput have motivated the continued development of different mass analyzer architectures, technologies and instrument configurations capable of balancing these parameters within practical instrument footprints ^7–12^. As a result, advances in time-of-flight (TOF) and other analyzer designs have remained an active area of research and development.

TOF mass analysers combine fast acquisition with full mass-range detection and are commonly used in workflows involving rapid separations and MS imaging applications. This has become increasingly apparent with recent advances in liquid chromatography (LC) separation technology, where reductions in extra-column volume and improved column temperature management have resulted in significantly increased chromatographic peak capacity and analysis speed ^13,14^. However, in conventional TOF configurations, improvements in resolving power are typically achieved by increasing the physical flight path length, which can lead to larger instrument dimensions and/or reduced duty cycle and ion transmission ^15^.

Multi-reflecting time-of-flight (MRT) analyzers were developed to address these limitations by extending the effective ion flight path through repeated reflections while maintaining compact analyzer geometries ^7,10,15,16^. Recent studies have shown that compact MRT-based instruments can achieve high resolving power and mass measurement accuracy at acquisition speeds compatible with high-throughput operation ^17^, and can address dynamic range requirements, for example, for the detection of environmental pollutants in food safety analysis^18^. Further development of MRT mass spectrometer architecture and acquisition functionality remains of interest to increase performance and to support demanding qualitative and quantitative LC-MS based omics and other workflows.

## Experimental

### Samples

EquiSPLASH™ was obtained from Avanti Research (Alabaster, AL). NIST® SRM® 1950 was from National Institute of Standards and Technology (Gaithersburg, MD). Human sera corresponding to human subjects diagnosed with bipolar or schizophrenia, in addition to healthy controls were purchased from Innovative Research (Novi, MI). Normal human urine was obtained from Innovative Research as well. MassPREP™ *E. coli* Digest Standard was from Waters Corporation (Milford, MA). All other chemicals and standards were from Sigma-Aldrich (St. Louis, MO), unless stated otherwise.

### Sample Preparation

EquiSPLASH, consisting of a mixture of deuterated internal standards representing various lipid classes, was dissolved in isopropanol and further diluted to 0.01 µg/mL in 100 mM aqueous ammonium acetate and isopropanol (10:490, *v/v*) for infusion. Additionally, for LC-MS analysis, EquiSPLASH was diluted in isopropanol to 0.1 µg/mL and in NIST SRM 1950 to 1 µg/mL.

Lipid extracts were prepared as previously described ^19^. In short, 100 µL serum was aliquoted and combined with three parts of isopropanol, which had been pre-cooled at −20 °C. Samples were then vortexed for 1 min before resting at room temperature for 10 minutes. To ensure efficient protein precipitation, samples were incubated at −20 °C overnight. Following incubation, samples were centrifuged at 14,000 g for 20 minutes. The resulting organic layer was removed and placed into fresh Eppendorf tubes. A pooled quality control (QC) sample was prepared by combing equal 50 µL aliquots of each serum sample prior to extraction with isopropanol to enable the quality of the analysis to be monitored ^20,21^. Prior to LC-MS analysis, the water content of each extract was adjusted to 50% with 50 µL of each sample being transferred to LC-MS total recovery vials (Waters Corporation).

Quantitative calibration standards were prepared by combining individual stock solutions of codeine, dihydrocodeine, fentanyl, oxycodone, and 6-acetylmorphine at 500 µg/mL each, representing a structurally diverse group of opioid analgesics and a specific heroin biomarker, respectively, followed by serial dilution in urine matrix to generate a calibration range of 0.056 to 10,000 ng/mL. Each standard was then further diluted by combining 100 µL of each standard individually with 400 µL of water in LC-MS maximum recovery vials MassPREP *E*.*coli* Digest Standard was dissolved in a 5% aqueous acetonitrile solution and diluted to a final concentration of 1 µg/µL in 0.1% aqueous formic acid solution.

### Infusion

MS and MS/MS direct infusion experiments were conducted with a syringe pump operated at a constant flow rate of 5 µL/min using 100 mM aqueous ammonium acetate and isopropanol (10:490, *v/v*) as the solvent system.

### Liquid Chromatography

Chromatographic small molecule separations were conducted with an ACQUITY™ Premier UPLC™ System (Waters Corporation). For the lipid standard analysis and small molecule omics discovery experiments, the solvents consisted of mobile phase A, comprising acetonitrile, 0.1% aqueous formic acid solution, and 10 mM aqueous ammonium formate (600:390:10, *v/v/v*), and mobile phase B, comprising isopropanol, acetonitrile, and 10 mM aqueous ammonium formate (900:90:10, *v/v/v*). A 4.5 min gradient was implemented using an ACQUITY Premier CSH™ C18, 1.7 μm, 1 x 50 mm Column (Waters Corporation), starting with a solvent composition of 50% mobile phase B. Over 2.3 minutes, mobile phase B was ramped to 80% before reaching 99% at 3 min. The composition was held at 99% mobile phase B for 0.3 minutes and re-equilibrated to initial conditions for 1.2 minutes. The column flow rate equalled 0.15 mL/min and the temperature was maintained at 55 °C.

For the quantitative small molecule drug analysis, the calibration standards were loaded on to an ACQUITY HSS T3, 1.8 μm, 2.1 x 100 mm Column (Waters Corporation), which was maintained at a temperature of 45 °C and operated with a flow rate of 0.6 mL/min. Mobile phase A consisted of 0.1 % (v/v) formic acid in water and mobile phase B of 0.1 % (*v/v*) formic acid in acetonitrile. Compounds were separated using a linear gradient: 1% mobile phase B from 0 to 0.3 min, increased to 50% over 7 min, to 70% at 8 min, before reaching 99% B at 9 min, followed by re-equilibration to 1% B by 10 min.

Protein digest separations were conducted using the same chromatographic LC system; however, here, an ACQUITY UPLC CSH130 C18, 1.7 μm, 2.1 x 100 mm Column (Waters Corporation) was used together with mobile phase A consisting of 0.1 % (*v/v*) formic acid in water and mobile phase B of 0.1 % (*v/v*) formic acid in acetonitrile. The column temperature was maintained at 45°C and linear gradients from 3 to 40 % mobile phase B were applied over durations from 1 to 15 (20 min inject-to-inject cycle time) and 1 to 55 minutes (60 min inject-to-inject cycle time) at a flow rate of 0.25 mL/min. Column re-equilibration times were 5 min. Protein digest on-column amounts were 2 or 4 µg for the 20 min gradient experiments and 2 µg for the 60 min gradient experiments.

### Mass Spectrometer Design and Operation

The geometry of the Xevo™ MRT P10 Mass Spectrometer (Waters Corporation, Wilmslow, UK) is shown in **Figure 1**, highlighting in red the design modifications relative to the original compact multi-reflecting instrument design ^7^. The main hardware changes include an increased sampling cone orifice of 0.9 mm, located at the entrance to the first vacuum stage and forming an integral part of the ion source, which necessitated increased backing pump capacity (EV-SA30, Ebara Corporation, Tokyo, Japan), and a revised orthogonal acceleration (OA) exit aperture with a larger diameter of 22 mm. Aberration and ion focusing has been described in detail previously ^7^. The larger OA exit aperture did not require additional analyzer design modifications, as its contribution to the overall aberration is minimal ^7^. MS and MS/MS acquisition speeds have increased up to 100 and 200 Hz respectively, with the option to configure the number of MRT reflections to tune mass resolving power and sensitivity. Typically, eight reflections were applied for the discussed applications, unless stated otherwise.

**Figure 1.**
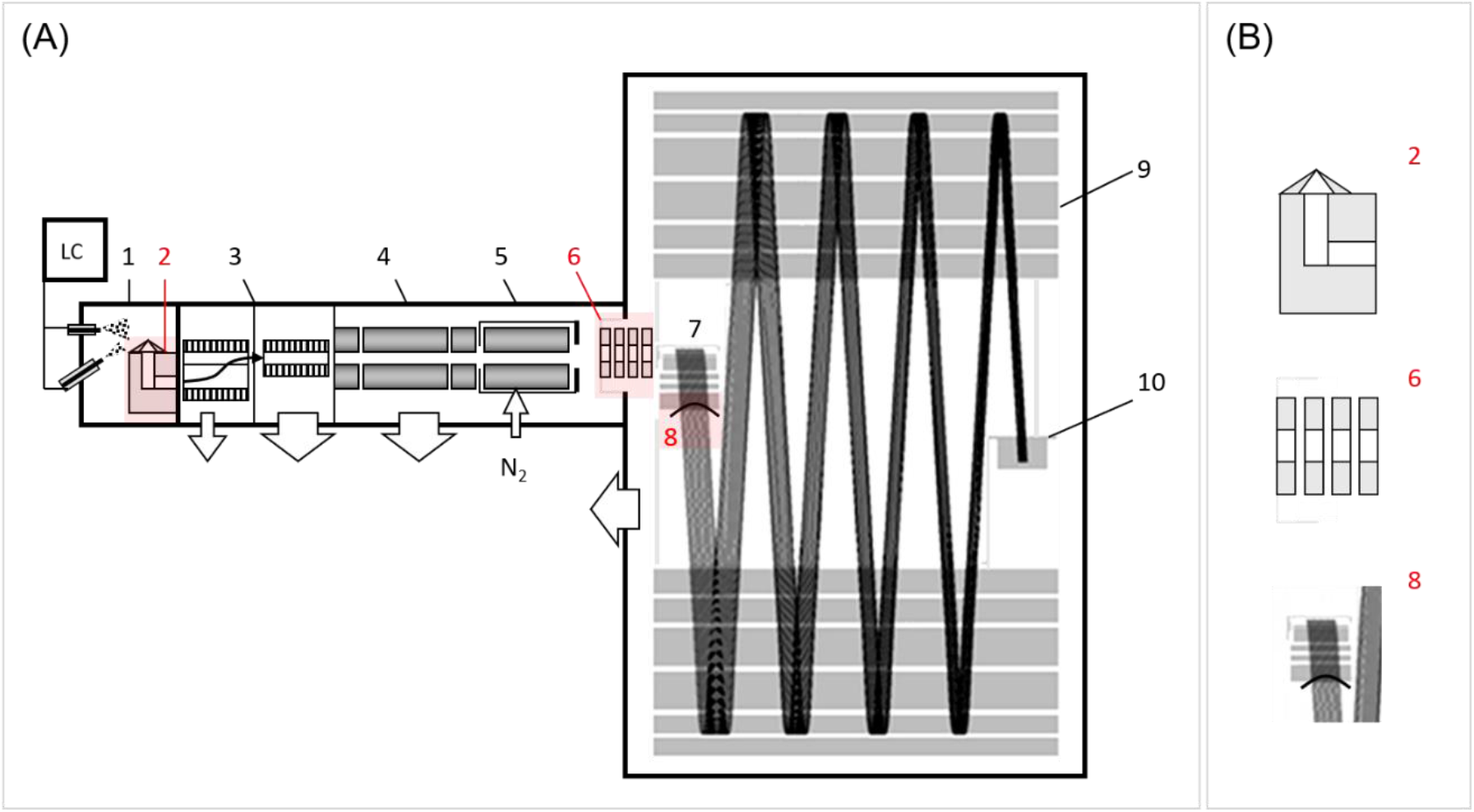
(A) Schematic quadrupole compact MRT instrument, consisting of the following main components, (1) ion source, (2) heated nozzle, (3) dual stage travelling wave ion guides (TWIGs), (4) analytical quadrupole, (5) collision cell, (6) lens system, (7) gridless orthogonal accelerator (OA) with a (8) trans-axial exit lens, (9) gridless ion mirrors, and (10) dual gain analogue-to-digital converter (ADC) detector. The white arrows show the location of the mechanical and turbo-molecular pumps. The main hardware modifications to the original design are highlighted in red and detailed in (B) include an increased 0.9 mm cone orifice residing in (1), and (8) a 22 mm OA exit aperture. Wide Band Enhanced Duty Cycle (WBEDC) functionality is incorporated within the lens system (6), located between the collision cell and the MRT analyser, and is discussed in detail in the Experimental and Results and Discussion sections.

Qualitative small molecule MS data were acquired using MS, MS/MS, and Data Independent Acquisition (DIA) in positive ionization mode. Infusion MS and LC-MS precursor-to-precursor MS/MS spectra were acquired with a low collision energy of 6 eV applied to the collision cell. For infusion MS/MS and LC-MS/MS experiments, spectra were collected using a 20 to 50 eV collision energy ramp. These conditions allowed assessment of spectral quality and mass resolution at different acquisition rates, ranging from 10 to 200 Hz. Small molecule data-independent acquisition (DIA) chromatograms and mass spectra experiments were typically acquired using a 22 to 55 eV collision energy ramp.

Quantitative small molecule LC-MS data were acquired using broadband DIA (MSE) ^22–24^ data acquisition in positive ionization mode. Precursor and product ion collision-induced dissociation (CID) spectra were collected from 50 to 1200 *m/z* at a scan rate of 10 Hz, with a collision energy of 6 eV applied to the collision cell for precursor ion data and a collision energy ramp from 20 to 45 eV for product ion CID data.

Small molecule discovery omics and protein digest MS data were obtained using a stepped quadrupole DIA acquisition method, referred to here as SONAR Pulse. This approach was conceptually described by Venable *et al*. ^25^ and variants ^26–28^ introduced later, followed by many adoptions, including scanning quadrupole ^29,30^ and ion mobility synchronized based approaches ^31^. Quadrupole isolation widths were 5 Da for all gradient duration and protein digest loading experiments. The MS1 and MS2 scan rates were 10 and 50 Hz, respectively, and the scan ranges 400 to 900 m/z and 150 to 2000 *m/z*. MS1 DIA chromatograms and spectra were collected at 6 eV and MS2 chromatograms and spectra using low 20 to 30 eV and high 50 to 60 eV collision energy ramps. Identical stepping quadrupole MS conditions were applied for the small molecule discovery omics experiments, using 22 to 55 eV collision energy ramps.

### Wideband Enhancement

Accumulating ions prior to orthogonal acceleration has been shown to significantly enhance the duty cycle on quadrupole time-of-flight mass spectrometers ^32,33^. This enhanced duty cycle (EDC) mode is achieved by synchronizing the pusher with the arrival of fragment ions released in a short pulse of 10-20 µs from the gas cell over a relatively narrow *m/z* range and has typically been used for targeted high-resolution analyses. In the method described here, ions are accumulated at the entrance to the lens system, component 6 in **Figure 1**, immediately downstream of the collision cell and prior to orthogonal acceleration. Following accumulation, ions are periodically released as ion packets and their arrival synchronized with the orthogonal acceleration sampling pulse by application of an appropriate delay time. Unlike Zeno-based approaches, which improve duty cycle through temporary ion trapping prior to TOF analysis ^34^, WBEDC enhances duty cycle by synchronizing the orthogonal acceleration sampling pulse with temporally broadened ion packets released from the collision cell, without the use of an ion trap.

By increasing the width of the pulse of ions released from the gas cell to 40-50 µs, a broader *m/z* range can be sampled by the orthogonal accelerator, but with concomitantly lower gains in duty cycle, which is referred to as wideband enhanced duty cycle (WBEDC). The net signal gain obtained with WBEDC and the other instrument modifications is illustrated in **Figure 2**. The contributions of the individual changes are specific to experimental conditions such as sample introduction flow rate, compound, and/or *m/z* range, and therefore cannot be represented by single gain values. **Figure 2** presents the MS/MS spectra of [Glu1]-Fibrinogen peptide B with amino acid sequence EGVNDNEEGFFSAR and elemental composition C_66_H_95_N_19_O_26_, infused at a concentration of 10^-7^ M, with and without WBEDC. The inset spectrum in the lower panel of **Figure 2** is shown on an expanded abundance scale, demonstrating that all product ions observed in the WBEDC data were also detected in the non-WBEDC experiment.

**Figure 2.**
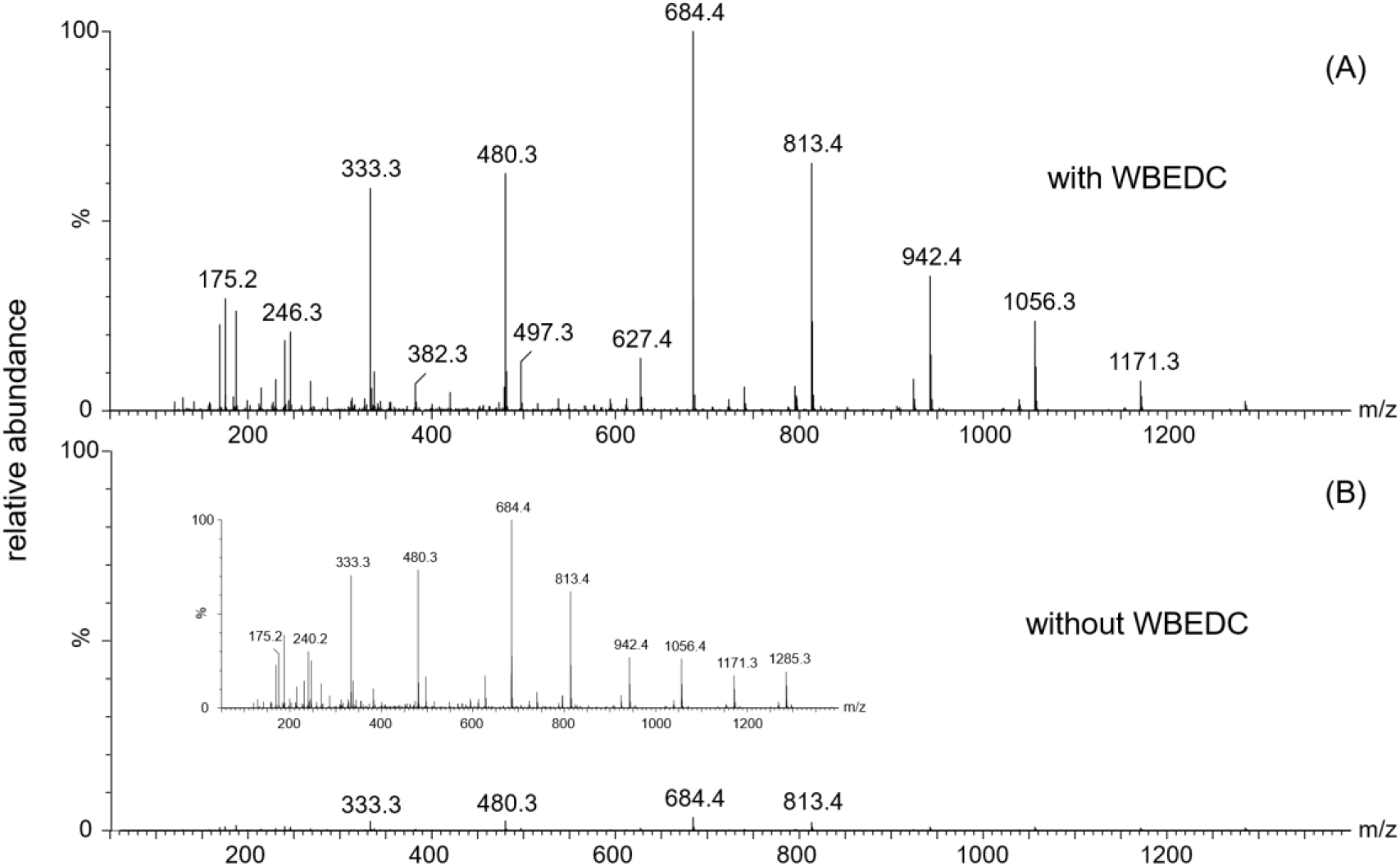
Collision-induced dissociation (CID) fragment ion MS/MS spectra of the doubly charged precursor *m/z* 785.84206 of [Glu1]-fibrinopeptide B. (A) With wideband enhanced duty cycle (WBEDC) applied, including hardware modifications as summarized in the **Experimental** section, and (B) without WBEDC. Shown as an inset in panel (B) is the non-linked abundance axis MS/MS spectrum of [Glu1]-fibrinopeptide B, indicating that all product ions were also observed in the non-WBEDC data.

### Data Acquisition and Processing

Data were collected and processed using waters_connect™ Software Platform v4.4 (Waters Corporation). Additional analysis of the data was achieved through DATA Convert (Waters Corporation) export into mzML format ^35^, which automatically determines and writes asymmetric MS1 isolation window offsets based on the transmission profile of the quadrupole analyzer, as these were found to be important for DIA acquisition methods that involved quadrupole isolation.

Small molecule lipidomics data were processed with MS-DIAL^36^ and protein digest data with DIA-NN v2.3 ^37^ (Aptila Biotech GmbH, Berlin Germany) or PEAKS Studio v13.1 ^38^ (Bioinformatics Solution Inc., Waterloo, Ontario, Canada). MS-DIAL searches were conducted with public MSP spectral databases (https://systemsomicslab.github.io/compms/msdial/main.html#MSP), with default identification, feature alignment based on retention time and accurate mass matching, and statistical analysis settings ^36^. Qualitative protein searches were conducted with a species specific FASTA amino acid sequence protein database (https://www.uniprot.org/proteomes/UP000000625), using fixed carbamidomethylation cysteine modification. The global protein group false discovery rate (FDR) was set at 1% with search tolerances determined automatically. All other settings were default.

Results and data visualization was performed using NumPy, Pandas, Matplotlib, and pyOpenMS Python™ libraries. Quantitative analysis was conducted with either waters_connect Software or Skyline v26.1 ^39^.

## Results

Various analyte mixtures were prepared to reproduce common omics sample types that are analyzed by qualitative and quantitative discovery MS experiments, including metabolites ranging from hydrophilic, low molecular weight compounds to more hydrophobic species such as lipids, as well as tryptic peptides. A series of analytical experiments were designed to assess the effect of instrument parameters, as far as possible in isolation, using specific acquisition methods. In addition to these basic performance tests, which primarily employed simple mixtures, more complex sample extracts and enzymatic hydrolysates were analyzed to assess performance from an application perspective, where the interplay of instrument parameters such as speed, selectivity, and specificity can have a marked effect on the outcome of an experiment.

### Acquisition speed

The effect of enhanced MS1 and MS2 acquisition speed on resolution and sensitivity was investigated using full-scan MS, MS/MS without collision energy applied to the collision cell, and CID MS/MS acquisition methods. Measurements were performed on an EquiSPLASH standard mixture at concentrations ranging from 0.01 to 1 µg/mL under infusion MS and LC-MS conditions, respectively. Infusion experiments were conducted using solvent standards. For the LC-MS experiments, samples were analyzed in the absence and presence of matrix, where the matrix consisted of SRM 1950, representing normal human plasma. Spectra were combined over a two-minute period for infusion data or across the chromatographic peak for LC-MS and LC-MS/MS data. Mass spectra were acquired using ten MRT reflections to achieve optimal mass accuracy and resolution. For the infusion experiments, an isotope pattern parameter (detector calibration threshold) of 0.1 ions per push was used. This threshold was selected to maintain operation within the linear response range of the detector and to ensure that the observed isotope patterns remained consistent with the expected isotopic distributions, thereby enabling reliable FWHM resolution measurements. The full width at half maximum (FWHM) resolution was largely independent of acquisition speed, as shown in **Figure 3** and **Supplementary Figure S1**, and summarized in **Supplementary Tables S1** to **S3**, with no significant change in resolving power at higher scan speeds. Although a small trend towards lower resolution was observed for some low *m/z* product ions in the MS/MS experiments, the observed differences remained within the experimental variability of the measurements. Acquisition speed had no effect on the accuracy of the isotopic distributions observed in either the MS or MS/MS data. The LC-MS and LC-MS/MS results are presented as average, normalized data for clarity in **Supplementary Figure S2**. Some larger variation in mass resolution was observed compared to the more controlled infusion experiments. However, overall trends were consistent, and no clear dependence on compound, *m/z*, or LC-MS or LC-MS/MS acquisition rate was observed. The retention time reproducibility, averaged over all compounds and LC-MS and LC-MS/MS experiments, and in the absence and presence of matrix, equaled 0.5 %CV. These findings support the use of higher acquisition rates without compromising data quality, which is of paramount importance for high-throughput ^13,14^ and imaging ^40^ applications.

**Figure 3.**
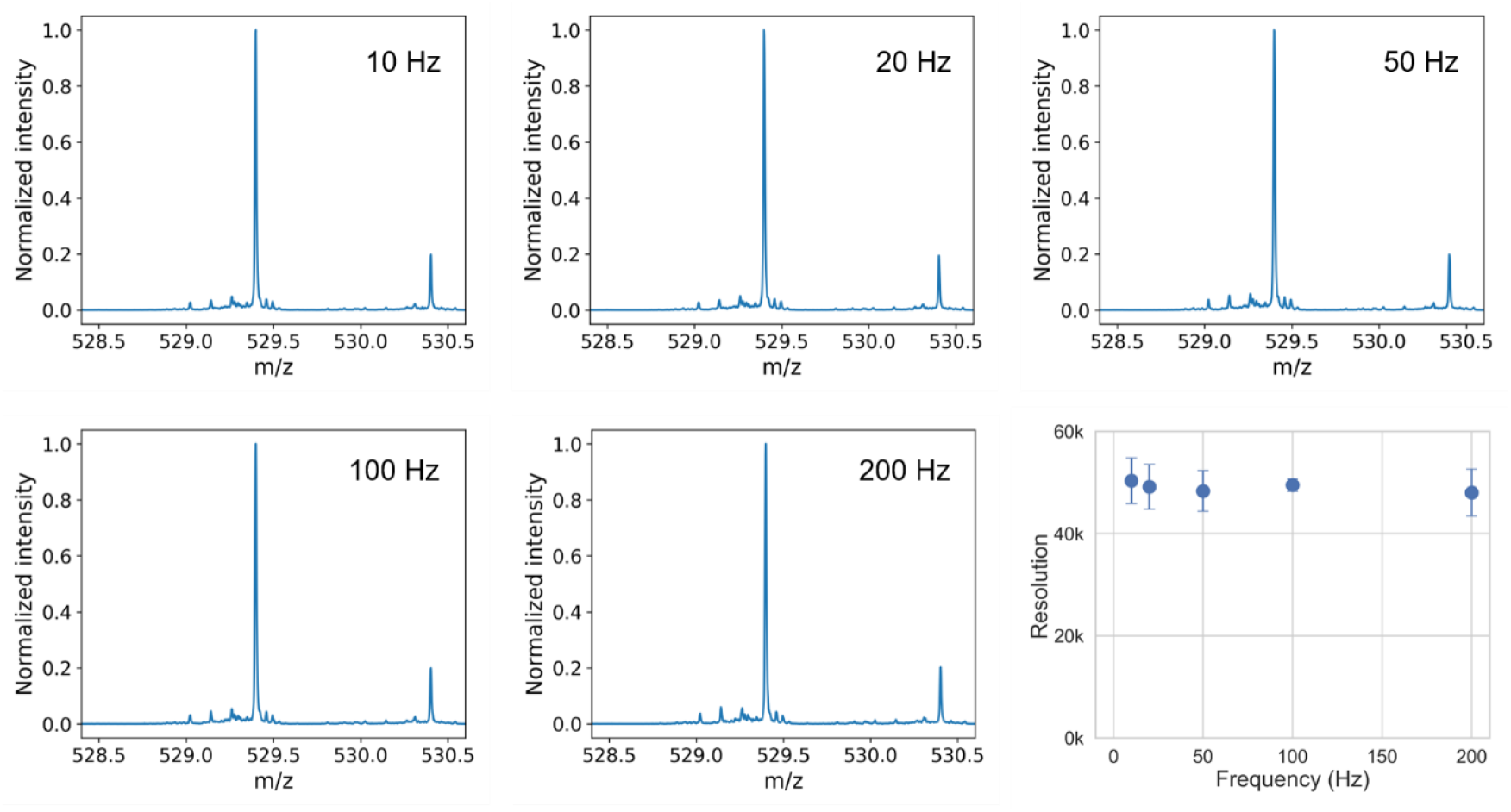
Summed detail infusion MS/MS spectra acquired following precursor ion isolation at *m/z* 529.3994 without CID (*m/z* 529.3994 > *m/z* 529.3994), collected at acquisition rates ranging from 10 to 200 Hz. FWHM resolution as function of MS/MS acquisition rate for 18:1(d7) Lyso PC (*n* = 6). Error bars represent two standard deviations.

### Quantitative Analysis

High-resolution and accurate-mass improve analytical specificity by resolving target analyte signals from background matrix components. This reduces interference and enhances signal-to-noise ratios, which is advantageous for monitoring and quantifying low-abundance metabolites or exogenous compounds in complex biological matrices such as urine. This approach was applied to the quantitative analysis of opioids and a drug-of-abuse marker using an unbiased broadband DIA (MSE) approach ^22–24^. This strategy enables quantitative measurements based on MS1, MS2, or a combination of both data levels. Representative Base Peak Intensity (BPI) and Extracted Ion Chromatograms (EICs) are shown in **Supplementary Figure S3**. These illustrate the complexity of the samples and the performance of the applied reversed-phase chromatographic separation. The separation system achieved an estimated peak capacity of approximately 273, with an average peak width at half height W_0.5_ of 1.7 s. Data acquisition was performed at 10 Hz, resulting in more than 20 data points per chromatographic peak per MS level, or over 40 points per chromatographic peak in total. These results demonstrate that the compact MRT mass spectrometer design supports acquisition rates suitable for high-throughput chromatographic separations while maintaining robust quantitative performance.

Extraction of specific accurate-mass MS1 precursor ions and selected broadband DIA MS2 product ions was used for quantification and confirmation, allowing the assessment of the quantitative performance and mass accuracy. The quantitative results, including the coefficient of determination (R^2^) obtained from the linear regression of response as a function of concentration, the lower limit of quantification (LLOQ), upper limit of quantification (ULOQ), average coefficient of variation (%CV), and linear dynamic range, are summarized in **Figure 4** and **Table 1**. Calibration levels were screened for acceptable precision (CV ≤ 20%) and proportional increases in mean response. The largest consecutive concentration range that met these criteria was selected for calibration, and an unweighted ordinary least squares (OLS) linear regression model was fitted to the accepted range. The LLOQ and ULOQ were defined as the lower and upper boundaries of this range. For clarity, only concentration levels included in the accepted calibration range are shown in **Figure 4**. R^2^ values were typically larger than 0.99, and the majority of the %CV values below 5% for the accepted concentration levels, without the use of internal standards, providing approximately 3.5 to 4.5 orders of linear dynamic range at the MS1 level and 2 to 3.5 orders at the MS2 level, which is favorable compared to previously reported metabolite quantitation using similar chemical compounds, matrix and acquisition method ^41^. Interestingly, 6-acetylmorphine behaved differently compared to the other four compounds. Its precursor *m/z* was affected by matrix interference at the MS1 level, and its MS2 fragment ions were produced less efficiently compared to those of the other compounds spiked into matrix. This is illustrated in **Supplementary Figure S4**, where both effects are evident as a relatively high MS1 mass error and the absence of detectable MS2 fragment ions at 4.57 ng/mL, respectively. In contrast, for dihydrocodeine, one of the other four compounds, the MS1 and MS2 detections at 4.57 ng/mL and 370.37 ng/mL were consistent in terms of mass accuracy and the abundance of the pseudo quantifier and qualifier ions relative to each other and to the precursor ion. Ultimately, however, signal-to-noise (S/N) determines the achievable lower limits of detection (LLOD) and quantification (LLOQ). These limits can be significantly improved, for example, by incorporating quadrupole isolation in combination with the previously described EDC approach. This type of MS/MS acquisition method, so called ToF MRM, is a targeted acquisition strategy that enables optimization, and provides a quantitatively viable alternative to conventional tandem quadrupole MS for targeted small-molecule analysis ^42^, but with enhanced analytical specificity.

**Table 1.** Quantitative summary MS1 and MS2 (pseudo quantifier) based quantification unbiased broadband DIA (MSE) for analgesics (opioids and a drug-of-abuse marker) in urine matrix; n = 5 (per concentration level).

| compound | $t_R$<br>(min) | adduct | MS1 | | | | | |
| --- | --- | --- | --- | --- | --- | --- | --- | --- |
| | | | $m/z$ (Da) | $R^2$ # | LLOQ<br>(ng/mL) | ULOQ<br>(ng/mL) | CV *<br>(%) | dynamic<br>range<br>(orders) |
| dihydrocodeine | 2.33 | [M+H]<sup>+</sup> | 302.1751 | 0.9983 | 0.056 | 3,333 | 4.5 | 4.8 |
| codeine | 2.37 | [M+H]<sup>+</sup> | 300.1543 | 0.9983 | 0.169 | 3,333 | 4.4 | 4.3 |
| oxycodone | 2.58 | [M+H]<sup>+</sup> | 316.1543 | 0.9998 | 0.500 | 10,000 | 3.4 | 4.3 |
| 6-acetylmorphine | 2.66 | [M+H]<sup>+</sup> | 328.1543 | 0.9965 | 1.520 | 10,000 | 3.5 | 3.8 |
| fentanyl | 4.79 | [M+H]<sup>+</sup> | 337.2274 | 0.9968 | 0.169 | 3,333 | 4.2 | 4.3 |

| compound | $t_R$<br>(min) | adduct <sup>Δ</sup> | MS2 | | | | | |
| --- | --- | --- | --- | --- | --- | --- | --- | --- |
| | | | $m/z$ (Da) | $R^2$ # | LLOQ<br>(ng/mL) | ULOQ<br>(ng/mL) | CV *<br>(%) | dynamic<br>range<br>(orders) |
| dihydrocodeine | 2.33 | QT [M+H]<sup>+</sup> | 227.1067 | 0.9966 | 4.57 | 3,333 | 5.8 | 2.9 |
|  |  | QL [M+H]<sup>+</sup> | 245.1172 |  |  |  |  |  |
| codeine | 2.37 | QT [M+H]<sup>+</sup> | 215.1067 | 0.9955 | 1.52 | 3,333 | 4.0 | 3.3 |
|  |  | QL [M+H]<sup>+</sup> | 282.1489 |  |  |  |  |  |
| oxycodone | 2.58 | QT [M+H]<sup>+</sup> | 298.1438 | 0.9993 | 4.57 | 10,000 | 4.9 | 3.3 |
|  |  | QL [M+H]<sup>+</sup> | 256.1332 |  |  |  |  |  |
| 6-acetylmorphine | 2.66 | QT [M+H]<sup>+</sup> | 268.1332 | 0.9991 | 123.0 | 10,000 | 6.8 | 1.9 |
|  |  | QL [M+H]<sup>+</sup> | 225.0910 |  |  |  |  |  |
| fentanyl | 4.79 | QT [M]<sup>+</sup> | 188.1434 | 0.9993 | 4.57 | 3,333 | 4.6 | 2.9 |
|  |  | QL [M]<sup>+</sup> | 105.0699 |  |  |  |  |  |

\* average coefficient of variation (CV) range for all concentration levels within the accepted linear dynamic range of the LC-MS experiments
<sup>Δ</sup> QT = quantifier product ion; QL = qualifier product ion

**Figure 4.**
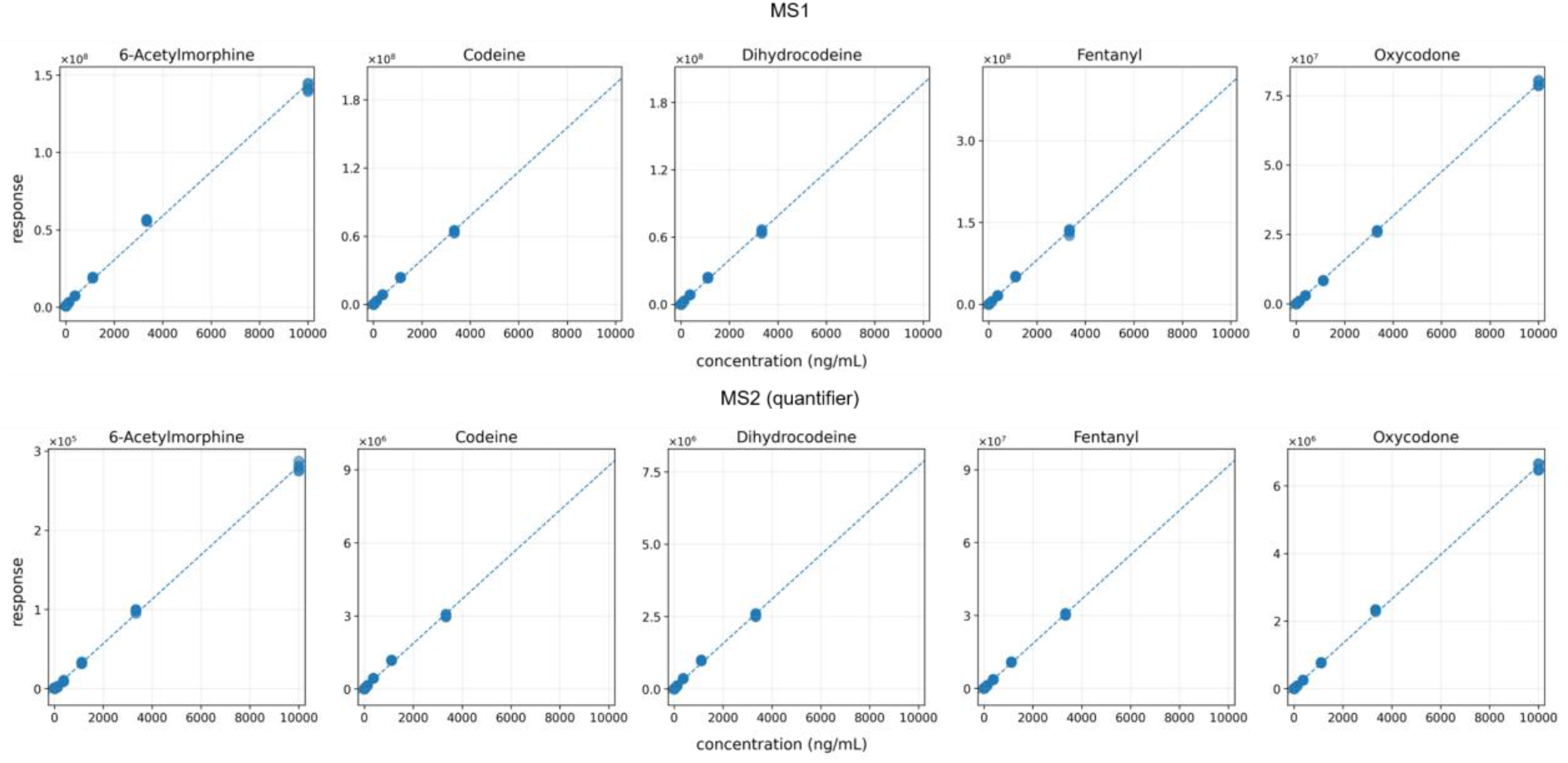
Quantitative broadband MS1 and pseudo quantifier MS2 DIA responses for analgesics (opioids and a drug-of-abuse marker) in urine matrix; n = 5 (per concentration level).

### Data Independent Acquisition Discovery Small Molecule Omics

The performance of stepped quadrupole DIA, named ‘*SONAR Pulse’*, was assessed and evaluated through the analysis of small molecule omics samples. This method combines a full-scan MS acquisition with rapid, sequential quadrupole isolation CID experiments, including WBEDC, across a defined *m/z* range using sequential mass selection windows, see **Experimental** section for method details. The experimental design for the lipid extract samples included three technical replicates of bipolar (*n = 5*) or schizophrenia (*n = 5*), in addition to healthy control (*n = 6*). Including six LC-MS conditioning runs and five study pool QC samples, which provided a total of 59 data files being analyzed. The samples were prepared as detailed in the **Experimental** section with the lipid extracts analyzed using a modified version of a previously published rapid microbore LC method ^43^. Prior to the analysis, conditioning injections with the pooled QC sample were performed to equilibrate the system and ensure column and instrumentation stability. Injections of the QC sample were interspersed throughout the sample batch after every tenth sample injection to enable consistency and QC checks. Representative base peak intensity (BPI) chromatograms of the four different sample types are provided in **Supplementary Figure S5**, demonstrating the achievable chromatographic separation efficiency under relatively fast chromatographic gradient conditions. While some minor differences in chromatographic BPI profiles can be observed, most of the differentiation is likely achieved through quantitative high-resolution mass spectrometric analysis of the samples.

MS-DIAL was employed for the analysis of the data, with a search example for one of the study pool QC samples shown in **Figure 5**. The precursor extracted ion chromatogram (EIC) is shown in **Figure 5A**, with the peak of interest highlighted blue and indicated by a red line. The corresponding putative identification demonstrates the type of sub-ppm mass accuracy and spectral MS2 clarity, shown in **Figure 5B** and **Figure 5C**, respectively, that can be obtained using stepped quadrupole DIA data with compact multi-reflecting ToF instrumentation and a fast LC method. The theoretical masses of the main fragment ions and corresponding mass errors for the annotated peaks in **Figure 5** are provided in **Supplementary Table S4**. This example also demonstrates the high specificity of the MS2 data, resulting in data-dependent acquisition (DDA) quality spectra. The deconvoluted EICs shown in **Figure 5D** illustrate the product ion information that can be obtained using this workflow. Increased MS/MS sensitivity improves the quality of spectral deconvolution and thereby increases confidence in compound identification. Moreover, more than 10 data points were obtained over the chromatographic peaks, affording sufficient peak sampling to enable quantitative precision for biomarker discovery studies. This is also reflected in the QC overview shown in **Figure 6A**, where the unsupervised principal component analysis (PCA) result on unfiltered, total ion current (TIC) normalized data indicates that *i)* the technical replicates clustered very tightly together, suggesting minimal experimental variation, *ii)* the study pool QC samples clustered together in the center of the scores distribution, indicating consistent data acquisition, and *iii)* separation of the control subject samples from the bipolar and schizophrenia subject samples, with partial separation of the bipolar and schizophrenia samples. These observations demonstrate the quality and consistency of the acquired lipidomics data and suggest the potential of the platform for detecting biologically relevant differences in lipidomic profiles.

**Figure 5.**
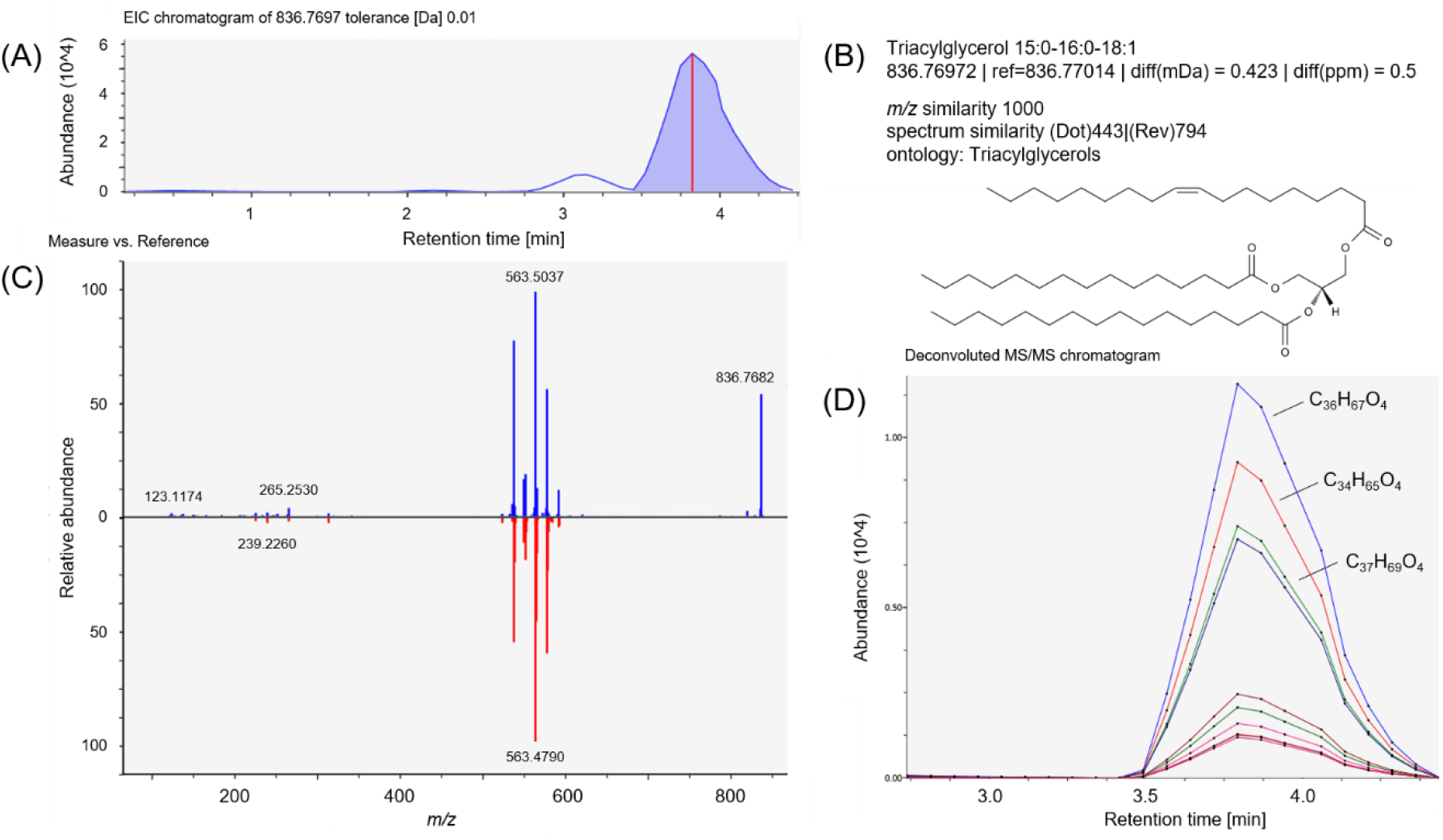
Qualitative Stepped Quadrupole DIA plasma lipid extract annotation showing the tentative identification of Triacylglycerol 15:0-16:0-18:1, comprising (A) precursor Extracted Ion Chromatogram (EIC), (B) search identification metrics, (C) measured (top) and experimental library reference spectrum (bottom) mirror spectra comparison, and (D) product ion EICs associated with the tentative identification following deconvolution.

**Figure 6.**
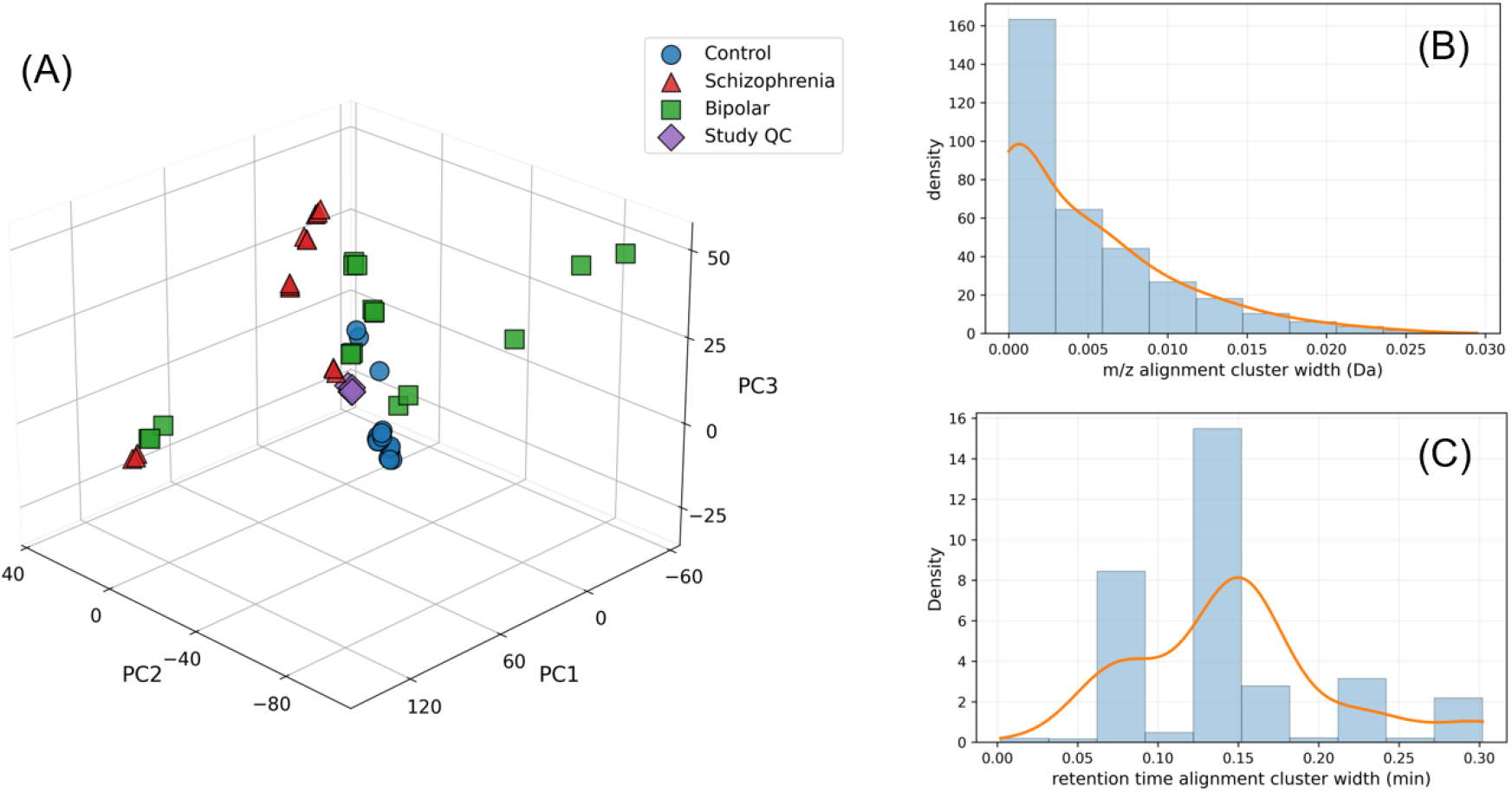
A) Unsupervised Principal Component Analysis (PCA) of unfiltered SONAR Pulse stepped quadrupole DIA plasma lipid extract LC-MS data. Alignment quality metrics showing the distributions of feature alignment cluster widths for (B) m/z and (C) retention time following retention time outlier filtering. The bars represent binned data and the curves fitted distributions.

The cluster widths afforded by the accuracy and precision of the mass measurement, and the retention time stability of the separation system, are shown **Figure 6B and Figure 6C**, respectively. As alignment quality metrics, *m/z* and retention time cluster widths were calculated and visualized. After excluding singleton clusters, retention time cluster width outliers were removed using the 1.5 times interquartile range (IQR) criterion, leaving 11,189 clusters for analysis. Cluster width was defined as the difference between the maximum and minimum *m/z* or retention time values within each cluster. The mean *m/z* cluster width was 4.746 mDa, with a standard deviation of 5.447 mDa, a median of 3.240 mDa, and an interquartile range (IQR) of 7.440 mDa. The mean retention time cluster width was 0.647 min, with a standard deviation of 0.668 min, a median of 0.152 min, and an interquartile range (IQR) of 1.131 min. The distribution of *m/z* alignment cluster widths provides an indication of the quality of mass resolution and accuracy. These data show an alignment precision of < 0.025 Da. However, a large proportion of the aligned features, about 84%, are < 0.01 Da and are therefore considered ultra-narrow width, typically associated with FT-ICR or Orbitrap mass analyzers ^44^. The LC separation domain also exhibits minimal run-to-run variation across samples, with all values < 0.3 min and the majority between 0.07 and 0.18 min. This high precision dramatically reduces the likelihood of false positives and decreases the chance of incorrectly grouping compounds. Minor alignment errors that could result in missing values are therefore minimized ^45^.

### Protein digest analysis

The analysis of proteolytic peptide digests using the compact MRT mass spectrometer was briefly evaluated, but a detailed description will be the subject of a separate publication. Peptide analysis is routinely applied in applications such as biopharmaceutical characterization, peptide bioanalysis and pharmacokinetics, and bottom-up proteomics. An example BPI chromatogram for the analysis of an *E. coli* tryptic digest together with representative SONAR Pulse DIA MS2 spectra from low-, mid-, and high-abundance proteins are shown in **Supplementary Figure S6**. Note the expected increasing apparent background, non-annotated signal, with decreased abundance. The number of identifiable and quantifiable proteins is influenced by various experimental parameters. In this study, gradient duration and on-column amount were varied, with the results summarized in **Supplementary Figure S7**, showing the number of detected and quantified proteins, as well as the experiment wide protein detection intersection, suggesting that maximum protein identification rates were already reached for the fastest gradient with the lowest amount of tryptic *E. coli* digest injected, which could be contributed to the limited complexity of this proteome. However, extending the gradient or increasing on-column load increased the number of detected and quantifiable peptides more significantly, as illustrated in **Supplementary Figure S8**. Although extending the gradient or increasing the on-column load resulted in only a modest increase in protein identifications, the corresponding increase in detected peptides provided greater amino acid sequence coverage, which could be important for certain applications or for the detection of low-level sequence variants or post-translational modifications. The highly reproducible observed quantifiable peptide detection results did not necessarily translate into similar reproducible protein quantification results, although still acceptable for most relative quantification studies, suggesting multiplication of error or variation when expressing protein CV values.

## Discussion and Conclusions

Given the increasing demand for higher sample throughput, high-resolution mass spectrometry-based omics workflows require continued improvements in resolving power, mass accuracy, and acquisition speed. These instrument parameters and characteristics are often interdependent; thus, the extent to which acquisition rate can be increased without affecting data quality remains an important practical analytical consideration. However, this is also directly influenced by instrumentation, and analyzer, sensitivity, as it ultimately determines signal intensity at a given acquisition rate. At the same time, the highest possible dynamic range should be retained. These core analytical instrument metrics, that is, speed, sensitivity, and selectivity, have been described and discussed in the context of enhancements applied to a compact MRT design.

The data and results presented here indicate that, by increasing MS/MS sensitivity, practical acquisition speeds can be increased up to 200 Hz without introducing significant losses in data quality, resolving power or isotopic ratio accuracy. The presence of matrix did not introduce significant variability, with the observed linear dynamic range spanning 3.5 to 4.5 orders of magnitude and lower limits of quantitation in a urine matrix at the sub-ng/mL level for analgesics. A SONAR Pulse DIA method was applied to the omics discovery analysis of lipid plasma extracts from mental disorder samples and an *E. coli* proteome standard, yielding data sets of sufficient quality for both qualitative identification and quantitative assessment under the applied acquisition conditions. These observations indicate that the acquisition method is compatible with demanding complex omics workflows requiring high acquisition rates combined with reproducible feature detection. Together, these results demonstrate that the combined hardware and acquisition advancements implemented in the system deliver improved sensitivity, higher acquisition rates, and enhanced quantitative performance, thereby extending the applicability to a range of demanding high-throughput and complex-matrix workflows and applications.

## Author Information

### Author Contributions

Conceptualization, JW, JPCV, JIL; Methodology, JW, SF, LAG, MED, MEP, RL; Writing – Original Draft Preparation, JPCV; Writing – Review and Editing, All authors.

### Notes

JW, SF, LAG, MED, MEP, RL, JPCV, and JIL are employed by Waters Corporation. ACQUITY, CSH, MassPREP, UPLC, Waters, waters_connect and Xevo are trademarks of Waters Corporation or its affiliates. Python is a registered trademark of the Python Software Foundation. SPLASH is a trademark of Avanti Polar Lipids, LLC. SRM is a registered trademark of NIST. US Disclaimer: For Research Use Only. Not for Use in Diagnostic Procedures. Rest of World Disclaimer: For Research Use Only

## Acknowledgement

The authors thank Xevo MRT Program and Mass Spectrometry Consulting for their support during the preparation of the manuscript. Mike McCullagh is acknowledged for proofreading the manuscript.

## Data Availability

The quantitative analysis small molecule LC-MS data have been deposited to the ProteomeXchange Consortium via the Panorama Public partner repository (https://panoramaweb.org/0gdDTL.url). All other data are available upon request.

