## supplemental for "Advances in the Design and Functionality of a Compact Multi-Reflecting Time-of-Flight Mass Spectrometer"

**Table S1.** Infusion MS resolution measurements (n = 6) for solvent standards.

| | | acquisition rate (Hz) | | | | |
|---|---|---|---|---|---|---|
| compound | *m/z* (Da) | 10 | 20 | 50 | 100 | 200 |
| 15:0-18:1(d7) DAG | 605.5844 | 55,336 | 54,465 | 55,977 | 61,481 | -- |
| 15:0-18:1(d7) PC | 753.6134 | 58,284 | 57,397 | 58,017 | 58,031 | -- |
| 15:0-18:1(d7) PE | 711.5664 | 53,041 | 51,783 | 53,223 | 53,189 | -- |
| 15:0-18:1(d7)-15:0 TAG | 829.7985 | 52,181 | 48,789 | 50,999 | 52,400 | -- |
| 18:1(d7) Lyso PC | 529.3994 | 58,840 | 56,467 | 57,604 | 55,463 | -- |
| 18:1(d7) Lyso PE | 487.3524 | 52,319 | 51,616 | 53,223 | 56,881 | -- |
| C15 Ceramide-d7 | 531.5477 | 52,177 | 50,461 | 53,950 | 53,951 | -- |
| d18:1-18:1(d9) SM | 738.6470 | 51,615 | 51,236 | 52,392 | 56,471 | -- |

**Table S2.** Infusion MS/MS resolution measurements (n = 6) for solvent standards (precursor *m/z* > precursor *m/z*).

| | | acquisition rate (Hz) | | | | |
|---|---|---|---|---|---|---|
| compound | *m/z* (Da) | 10 | 20 | 50 | 100 | 200 |
| 18:1(d7) Lyso PC | 529.3994 | 50,340 | 49,169 | 48,354 | 49,501 | 48,027 |

**Table S3.** Infusion MS/MS resolution measurements (n = 6) for solvent standards (precursor *m/z* > product *m/z*).

| | | acquisition rate (Hz) | | | | |
|---|---|---|---|---|---|---|
| compound | *m/z* (Da) | 10 | 20 | 50 | 100 | 200 |
| 18:1(d7) Lyso PC | 184.0733 | 45,931 | 44,651 | 45,311 | 42,053 | 40,199 |

**Table S4.** Theoretically predicted and experimentally detected fragment ion *m/z* values for TG 15:0/16:1/18:0.

| elemental composition fragment ion * | adduct | theoretical *m/z* Δ | observed *m/z* ‡ | error (ppm) | comment |
|---|---|---|---|---|---|
| $C_9H_{15}$ | [M+H]+ | 123.11737 | 123.1174 | 0.3 | low abundant |
| $C_{14}H_{29}$ | [M+H]+ | 197.22691 | -- | -- | |
| $C_{15}H_{29}O$ | [M+H]+ | 225.22183 | 225.2215 | -1.5 | low abundant |
| $C_{16}H_{31}O$ | [M+H]+ | 239.23748 | 239.2369 | -2.4 | low abundant |
| $C_{18}H_{33}O$ | [M+H]+ | 265.25313 | 265.2530 | -0.5 | |
| $C_{18}H_{35}O_2$ | [M+H]+ | 283.26369 | -- | -- | |
| $C_{36}H_{67}O_4$ | [M+H]+ | 563.50391 | 563.5037 | -0.4 | |
| $C_{52}H_{97}O_5$ | [M+H]+ | 801.73356 | -- | -- | |
| $C_{52}H_{99}O_6$ | [M+H]+ | 819.74413 | 819.7410 | -3.8 | low abundant |

* http://cfmid4.wishartlab.com/
Δ https://pnnl-comp-mass-spec.github.io/Molecular-Weight-Calculator-VB6/
‡ https://systemsomicslab.github.io/compms/msdial/main.html

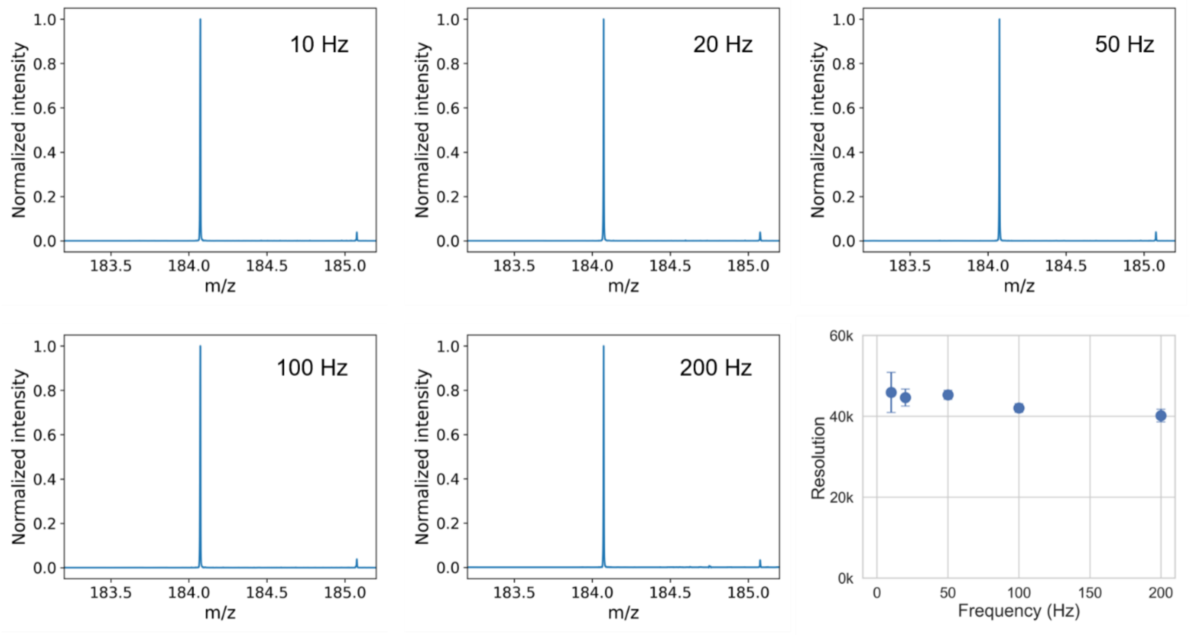


**Figure S1.** Summed detail infusion MS/MS spectra acquired following precursor ion isolation at *m/z* 529.3994 with CID (*m/z* 529.3994 > *m/z* 184.0733), collected at acquisition rates ranging from 10 to 200 Hz. FWHM resolution as a function of MS/MS acquisition rate for 18:1(d7) Lyso PC (*n* = 6). Error bars represent two standard deviations.

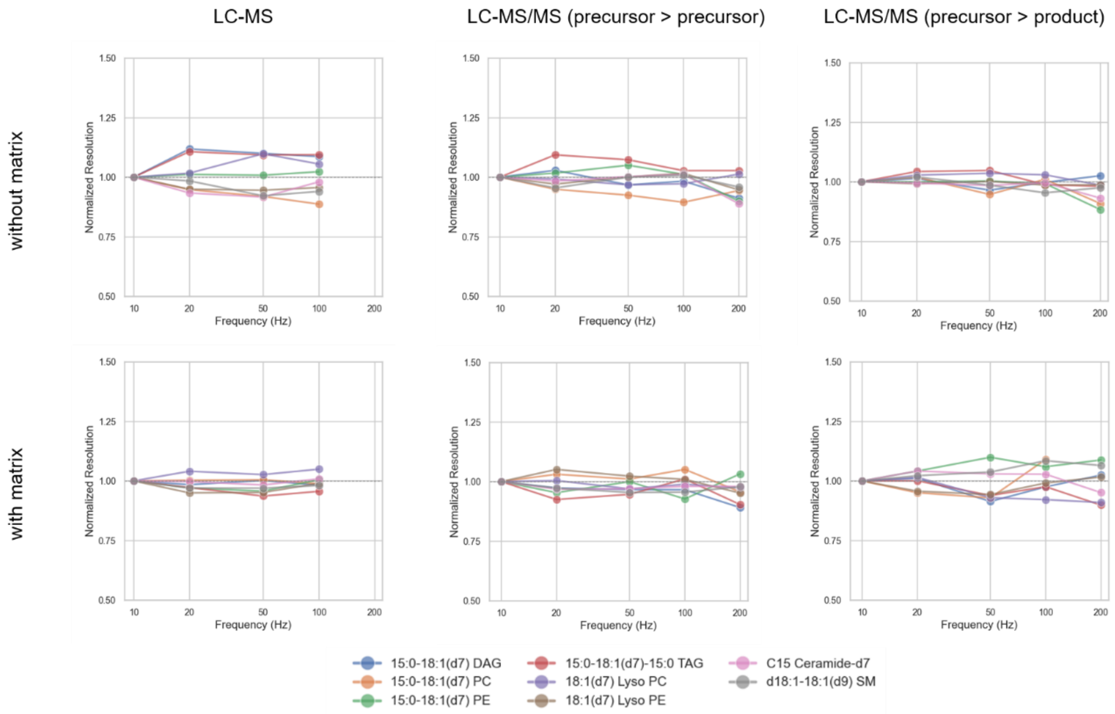


**Figure S2.** Normalized ($FHWH_{10\ Hz}$ = 1) average FWHM resolution (n = 6) as a function of LC-MS and LC-MS/MS acquisition rate in the absence (top row) and presence (bottom row) of urine matrix. Blue = 15:0-18:1(d7) DAG; orange = 15:0-18:1(d7) PC; green = 15:0-18:1(d7) PE; red = 15:0-18:1(d7)-15:0 TAG; purple = 18:1(d7) Lyso PC; brown = 18:1(d7) Lyso PE; pink = C15 Ceramide-d7; grey = d18:1-18:1(d9) SM. The LC-MS/MS matrix data were evaluated using the A+1 isotope peak.

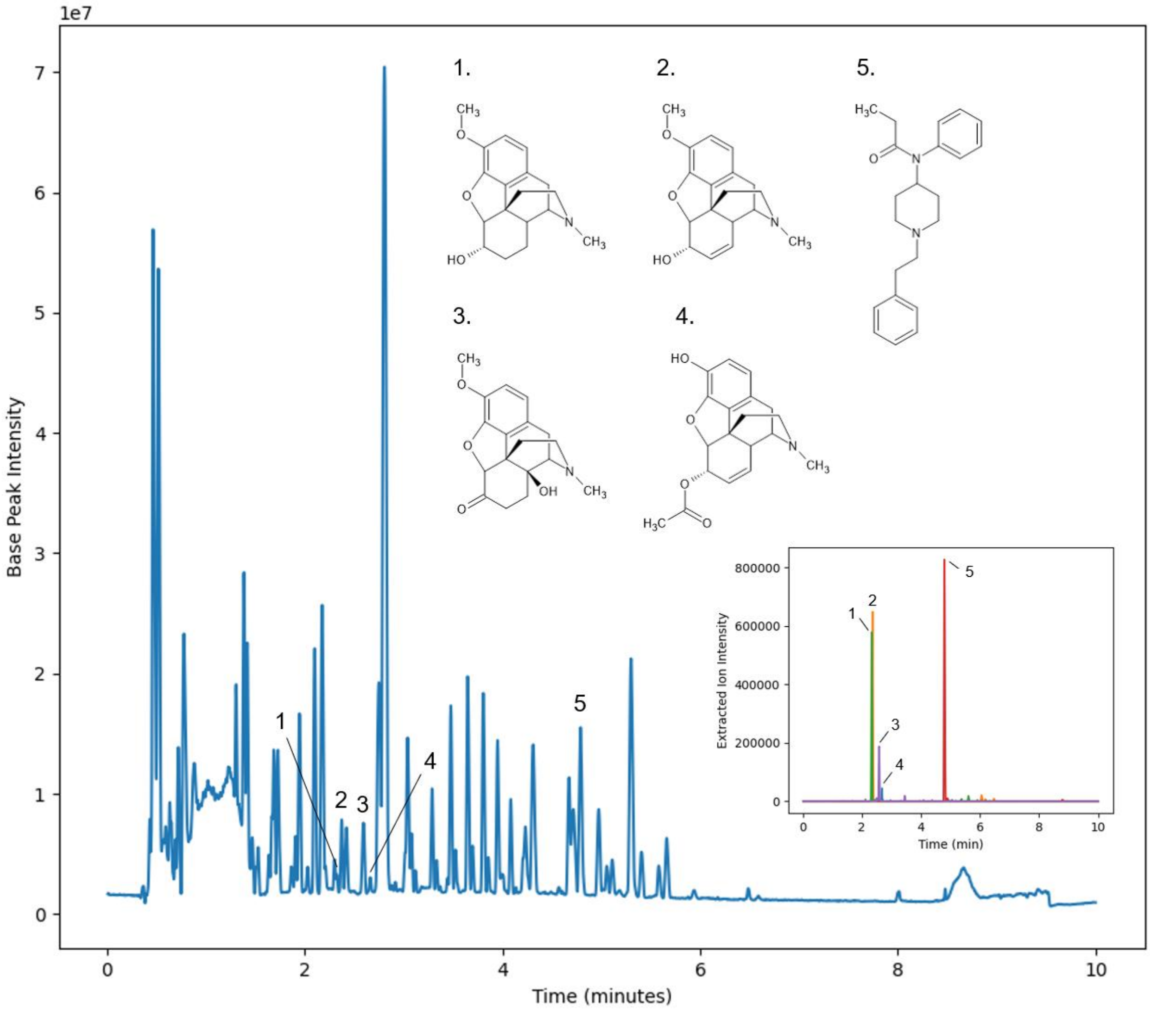


**Figure S3**. Base Peak Intensity (BPI) and precursor MS1 Extracted Ion Chromatogram (EIC), shown inset, broadband DIA (MSE) chromatograms of dihydrocodeine (1, green), codeine (2, orange), oxycodone (3, purple), 6-acetylmorphine (4, blue), and fentanyl (5, red), spiked at about 40 ng/mL into urine matrix.

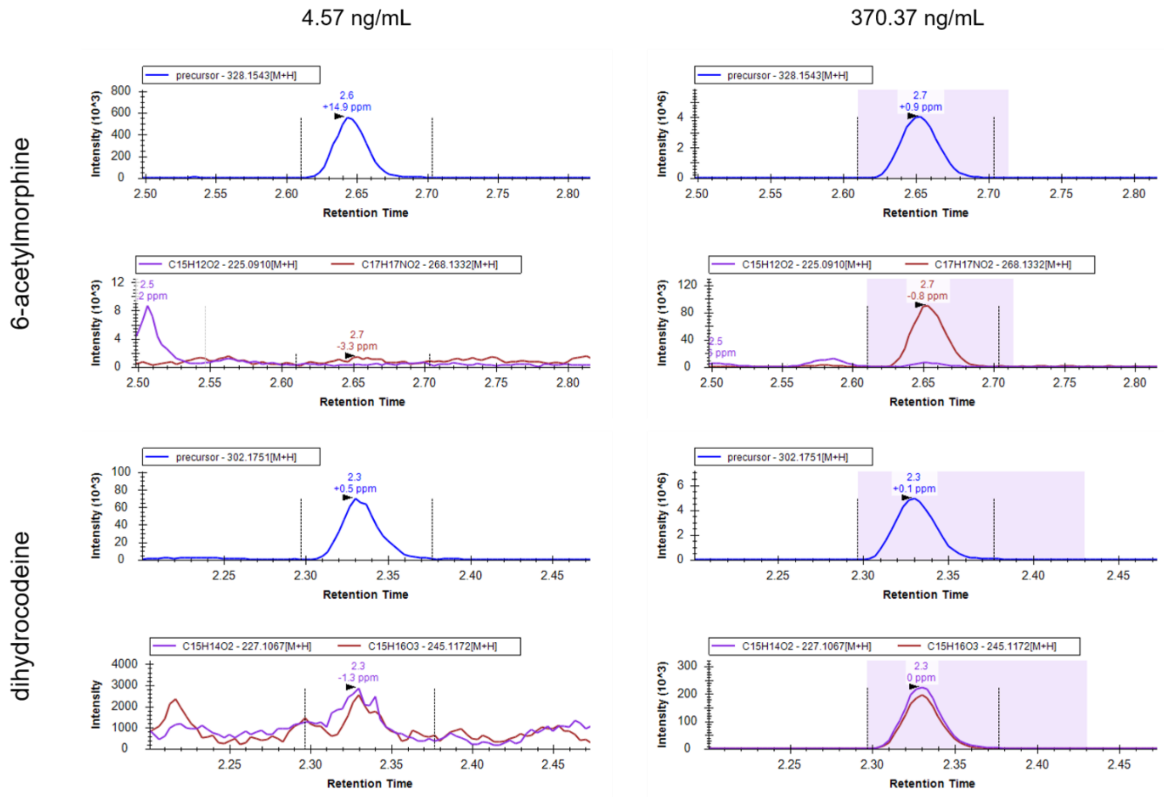


**Figure S4**. MS1 and MS2 Extracted Ion Chromatogram(s) (EIC) 6-acetylmorphine (top) and dihydrocodeine (bottom) spiked in urine matrix at 4.57 (left) and 370.37 (right) ng/mL.

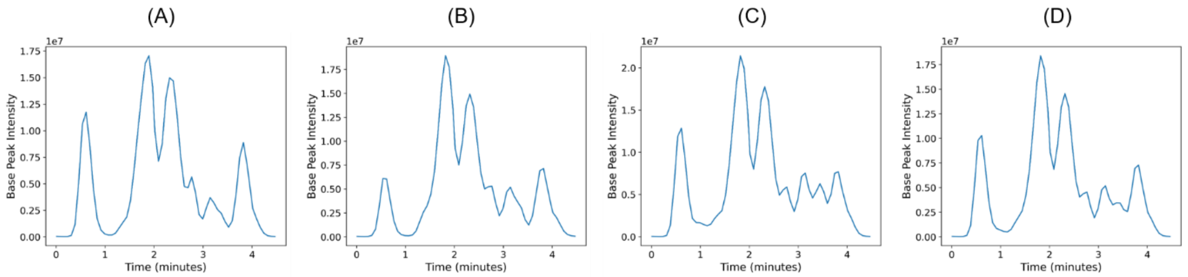


**Figure S5.** Example stepped quadrupole DIA Base Peak Intensity (BPI) chromatograms of (A) bipolar, (B) schizophrenia, (C) control, and (D) study pool QC lipid extract plasma samples from a mental disorder study using fast, narrow-bore LC separations.

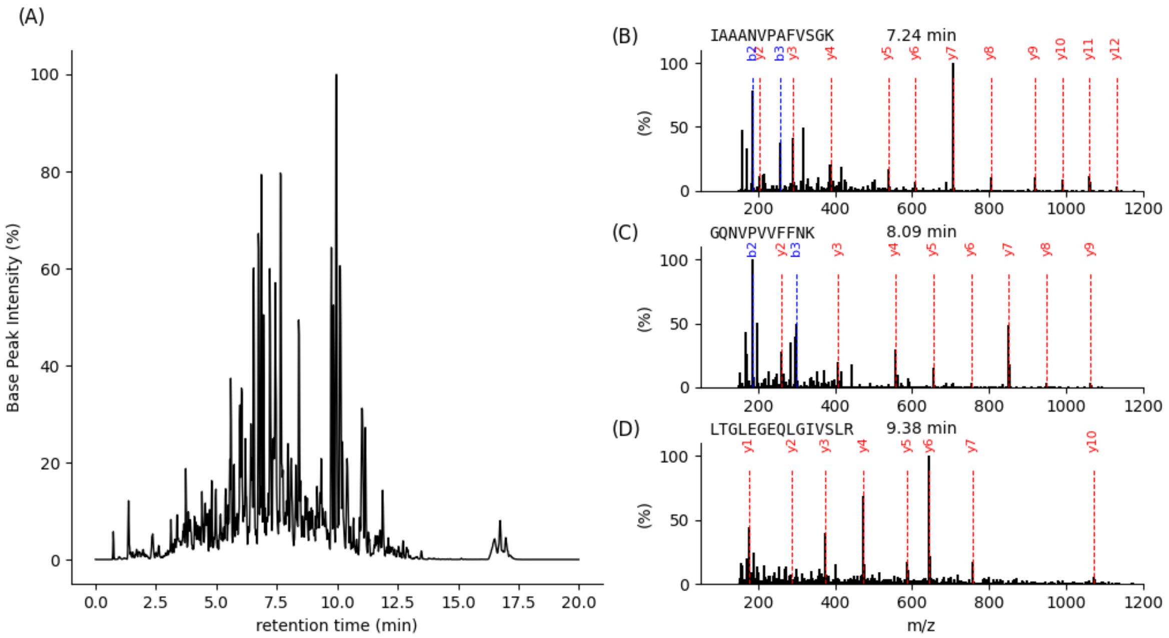


**Figure S6**. (A) Base-peak intensity (BPI) chromatogram and annotated raw MS2 product ion spectra showing *y*- and *b*-ions for (B) high-abundance (IAAANVPAFVSGK | P0ACF0 – DBHA_ECOLI | 7.24 min), (C) mid-abundance (GQNVPVVFFNK | P0AEE5 – MGLB_ECOLI | 8.09 min), and (D) low-abundance (LTGLEGEQLGIVSLR | P0A707 – IF3_ECOLI | 9.38 min) peptides acquired using stepped-quadrupole DIA with a 20 min gradient (inject-to-inject cycle time). Data were obtained from 4 µg of an *E. coli* tryptic digest loaded on-column, spanning at least two orders of magnitude in summed product ion area. Across the experiments, the root mean square (RMS) of the mean precursor and product ion mass errors was 0.47 and 2.45 ppm, respectively. The corresponding mean within-experiment standard deviations were 2.5 and 6.0 ppm.

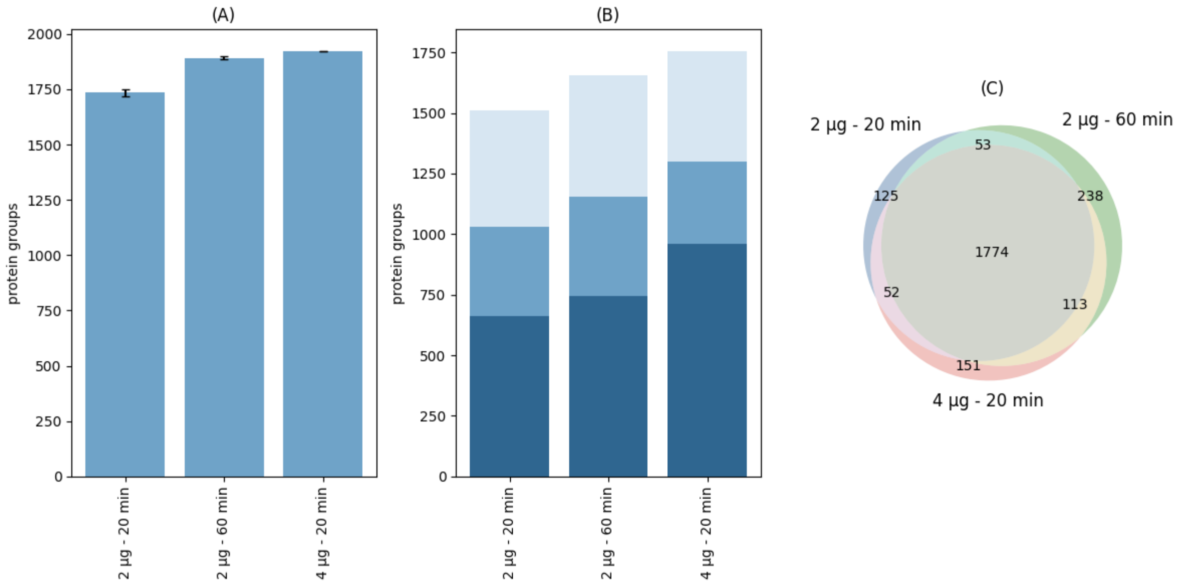


**Figure S7.** (A) Protein groups identified in an *E. coli* tryptic digest, (B) protein groups quantified with coefficients of variation (CV) below 10 % (dark blue), between 10 % and 20 % (mid blue grey), and above 20 % (light blue grey), and (C) protein group intersection for different on-column amounts and chromatographic gradient lengths (technical replicates, n = 3), determined using an individual-run-based analysis approach.

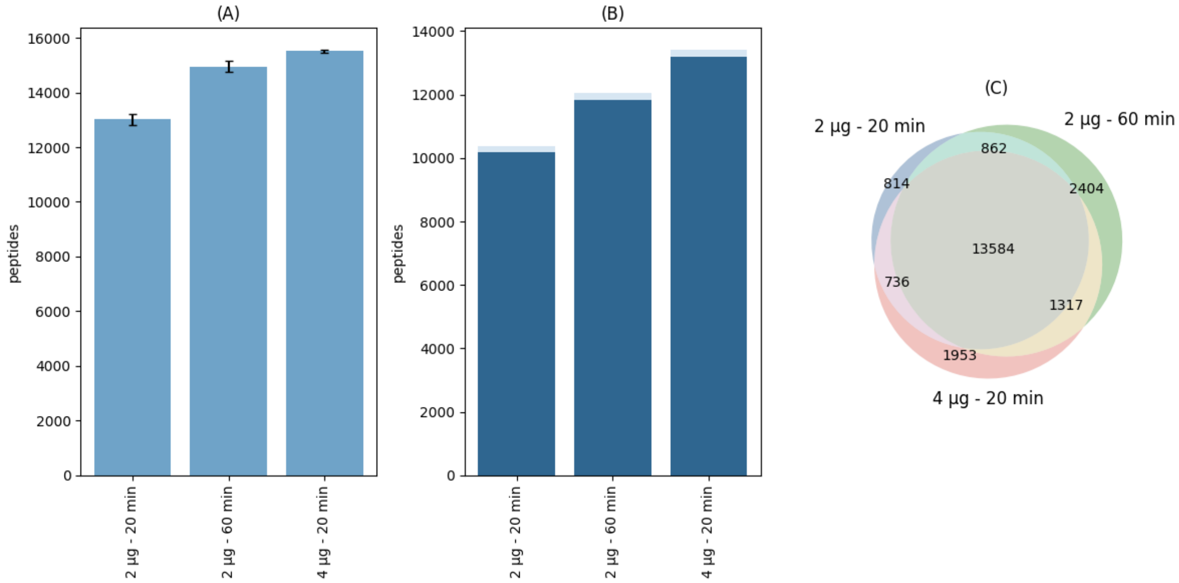


**Figure S8.** (A) Peptides detected in an *E. coli* tryptic digest, (B) peptides quantified with coefficients of variation (CV) below 10 % (dark blue), between 10 % and 20 % (mid blue), and above 20 % (light blue) and (C) peptide intersection for different on-column amounts and chromatographic gradient lengths (technical replicates, n = 3), determined using an individual-run-based analysis approach.
